# Screening of tomato seed bacterial endophytes for antifungal activity reveals lipopeptide producing *Bacillus siamensis* as a potential bio-control agent

**DOI:** 10.1101/2020.10.09.332627

**Authors:** Ayushi Sharma, Nutan Kaushik, Abhishek Sharma, Abhay Bajaj, Mandar Rasane, Yogesh S. Shouche, Takwa Marzouk, Naceur Djébali

## Abstract

The current study investigates the diversity pattern and fungicidal potential of bacterial endophytes isolated from two different organic varieties of tomato plants (V1 and V2). A total of seventy-four bacterial isolates identified by 16S rRNA sequencing revealed a single genus *Bacillus* with 16 different species. The Shannon diversity H’ (1.45), Simpson’s index of diversity (0.9), Magalef’ index (2.1), Evenness (0.96), and Species richness (8) indicated the high endophytic bacterial diversity in the V1 variety of the tomato. Bacterial endophytes isolated from both the varieties were screened for their antifungal activity against five economically critical fungal pathogens (viz., *Botrytis cinerea, Rhizoctonia solani, Fusarium solani, Verticillium lateritium*, and *Alternaria solani*) of tomato crop through dual culture assay. The data revealed *B. siamensis* KCTC 13613(T) as the most potent antagonist significantly (p < 0.05), inhibiting the mycelial growth between 75 to 90% against selected fungal pathogens. High bioactivity of lipopeptide extract of *B. siamensis* was recorded against *R. solani* with IC_50_ value of 72 ppm. The UPLC-HDMS analysis of this lipopeptide extract revealed the presence of, Surfactin and Bacillomycin D.

## INTRODUCTION

Tomato (*Solanum lycopersicum*) is a well-known vegetable crop due to its high nutritional values. Like many other crop plants, it suffers from various fungal diseases. Its property to bear a succulent fruit increases its susceptibility towards fungal attacks than other crop plants, which is an essential limiting factor in its production (Habiba et al., 2017). The key phytopathogens responsible for damaging this crop include *Rhizoctonia solani, Fusarium solani, Botrytis cinerea, Alternaria solani*, and *Verticillium sp*. Because of their diverse host spectra and soilborne existence, fungal phytopathogens are difficult to control (Lamichhane et al., 2017). The use of chemical fungicides is the most common strategy to prevent fungal pathogens (Windels and Brantner, 2005). However, due to the rising environmental contamination and appearance of the pathogen’s resistant races, seed bio priming with endophytes is being looked upon as an environmentally friendly option.

Endophytic bacteria play an essential task in managing plant health and diseases (Hazarika et al., 2019). These bacteria harbor inside the plant and contribute to reduced population densities of pathogens without stimulating hypersensitive reactions in the host (Hazarika et al., 2019; Roy et al., 2017). Bacterial endophyte composition varied among plants, organs, genotypes, tissues, cultivars, soil, and location (Kumar et al., 2020). The rhizosphere or phyllosphere work as a source for several endophytes; nevertheless, some bacterial species have been reported vertical transmission through seed (Truyens et al., 2015).

Many endophytic bacteria exhibit antagonistic ability towards fungal pathogens. *Bacillus* species produce heat and UV resistant spores that can withstand adverse environmental conditions, thereby becoming an attractive agent for commercial use in modern farming systems (Piggot and Hilbert 2004; Tiago et al., 2004). The antifungal ability of isolate *Bacillus subtilis* SCB-1 was identified against diverse fungal pathogens, including the *Alternaria* and *Fusarium* (Hazarika et al., 2019). Isolation and characterization of highly antagonistic *Bacillus* strains have reported volatile organic compounds against *Sclerotinia sclerotiorum* (Massawe et al., 2018).

In the present study, the diversity of the endophytic bacteria isolated from the various tissues of two different organic tomato varieties was evaluated. We also characterized the antifungal activity of an endophytic bacterial isolate, *Bacillus siamensis* KCTC 13613(T) and identified the major antifungal components through UPLC-HDMS analysis.

## MATERIALS AND METHODS

### Seed Collection

Two organic tomato varieties were used in this study for the isolation of bacterial endophytes. Both the varieties, i.e., Pusa Ruby (Maharashtra) (V1) and a local variety of Andhra Pradesh (Madanapalle) (V2) were procured from the online garden stores, Ugaoo and Organic Garten, respectively.

### Isolation of Bacterial Endophytic Strains

Surface sterilization of tomato seeds was performed to remove the epiphytic bacteria following the method described by Kumar et al. (2011). Seeds were first sterilized with 70 % ethanol for 2 minutes, followed by a 1 % sodium hypochlorite solution for 3 minutes. After that, surface-sterilized seeds were washed three times with autoclaved distilled water and dried with sterile blotting paper. For sterility check, imprints of dry surface-sterilized seeds were taken on Luria-Bertani agar medium. Seeds were then put for germination on sterile filter paper immersed with autoclaved distilled water in a petri dish at 27°C. For isolation, seedlings obtained after the nine days of germination were again surface sterilized with the method described above. After sterility check, each seedling was cut into different sections viz., root, hypocotyl, and cotyledon. Each part was further divided into various segments and placed on the Luria-Bertani agar plate. Plates were then incubated for 2-3 days at 27°C. Visually distinct bacterial colonies acquired from segmented seedlings were purified and maintained in LB agar slants/plates and glycerol stock at 4°C and −80°C, respectively.

### Identification of Bacterial Isolates and Construction of Phylogenetic Evolution

The identification of isolates was carried out at the Sequencing facility of National Centre for Microbial Resource (NCMR), National Centre for Cell Science, Pune.DNA extraction and purification was done using HiPurA™ 96 Bacterial Genomic DNA Purification Kit (Himedia), as per manufacturer’s protocol; followed by amplification of 16S rRNA gene using universal bacterial primers (27F,1492R). Amplified products were sequenced by Sanger method on ABI 3730xl Genetic Analyzer (Applied BioSystems). The sequences were aligned and evaluated for taxonomic identification by BLAST analysis (Boratyn et al., 2013). The phylogenetic tree was reconstructed by doing alignment using Clustal W and the evolutionary history inferred using the Neighbor-Joining method. A tree with 1000 bootstrap replicates was constructed using MEGA-X.

### Diversity Indices

Bacterial endophytes derived from organic tomato seedlings were grouped into their specific isolation sections, such as hypocotyl, root, and cotyledon, which facilitated the comparison between the isolates of the same or other variety. Species diversity was calculated using the Shannon diversity index to measure species evenness and richness (Chowdhary and Kaushik, 2017).

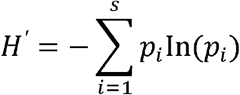

Where, *s* equals the number of species, and *pi* equals the ratio of individuals of species *i* divided by all individuals *N* of all species. The Shannon diversity index ranges typically from 1.5 to 3.5 and rarely reaches 4.5. Simpson’s index (D) was calculated to determine the dominance, the higher the value lower in the diversity (Ifo et al., 2016).

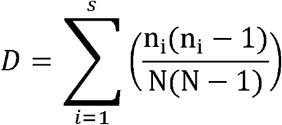

Where, *n_i_* is the number of individuals in the *i*^th^ species and TV equals the total number of individuals and Simpson’s index of diversity was calculated b

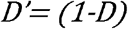

Other parameters, such as species evenness and richness, were also calculated (Ifo et al., 2016). Margalef’s index (*d*) also indicates the evenness (Kumar et al., 2006). A value for evenness approaching zero reflects large differences in the abundance of species, whereas an evenness of one means all species are equally abundant,

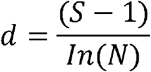

*S* is the total number of species; *N* is the number of individuals, and the natural logarithm.

To measure the similarity in the species composition for both varieties of tomato, we used Sorenson’s index of similarity using the equation,

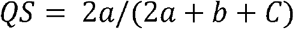

and Jaccard’s index of similarity using the equation,

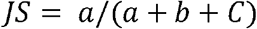

Whereas, ‘*a*’ denotes the number of bacterial species commonly shared by both the varieties, ‘*b*’ denotes the number of bacterial species found in V1, and ‘*c*’ denotes the number of bacterial species found in V2 (Chowdhary et al., 2015).

### In-vitro Antifungal Activity of Bacterial Endophytes

All the bacterial isolates were screened for their antagonistic activity against major pathogenic fungi of the tomato crop, namely*, Rhizoctonia solani* (ITCC-6430), *Fusarium solani* (ITCC-6731), *Botrytis cinerea* (ITCC-6011), *Alternaria solani* (ITCC-4632), and *Verticillium lateritium* (ITCC-2819) obtained from Indian Type Culture Collection (ITCC) at Indian Agricultural Research Institute (IARI), Pusa, New Delhi, India. Isolates were evaluated by dual culture assay on Potato Dextrose Agar (PDA) medium. Fully grown 7mm fungal disc was placed in the center of the PDA plate while bacterial isolate was streaked on both the sides of the fungal disc at equidistance. PDA plate inoculated only with the fungal disc was kept as control. After 3-5 days of incubation, plates were observed for the antagonism expressed by endophytic bacteria, and percentage growth inhibition was calculated. Growth inhibition (GI) was calculated as per the following:

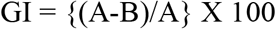

Where, A = radial growth of the plant pathogenic fungus in control; B = radial growth of the plant pathogenic fungus in the presence of endophytic bacterial strain (dual inoculation).

### Extraction and Purification of Lipopeptide

Bacterial endophyte, *B. siamensis*, with the most promising antagonistic activity against all the test pathogenic fungi, was further explored to produce antifungal lipopeptides. The lipopeptide extraction method involved acid precipitation and solvent extraction, as described by Romano et al. (2011). Briefly, extraction of lipopeptide from a cell-free supernatant was done by precipitation method at pH 2 using 6N HCl and incubated at 4°C overnight and then centrifuged at 12000 rpm for 15 minutes at 4°C. The pellet was extracted using a mixture of Chloroform: Methanol (2:1, v/v) followed by centrifugation for at 12000 rpm for 15 minutes at 4°C. The extract present in the supernatant was filtered and concentrated to dryness by rotary evaporation. Waters ACQUITY UPLC H-class with Synapt G2-Si High Definition Mass Spectrometry (HDMS) system with the C18 column was employed for lipopeptide profiling. The lipopeptide extract (10 mg) was dissolved in 10 mL of HPLC grade ethanol extracts, and a 10 μL sample was injected into UPLC coupled with HDMS.

### Antifungal Bioassay of Lipopeptide

The antifungal bioassay of lipopeptide was carried out by the agar diffusion method in PDA (Chowdhary and Kaushik, 2015). The lipopeptide was dissolved in ethanol to make a stock solution of 1000 ppm. From the stock solution 50 μL, 150 μL and 300 μL along with ethanol control were spot-inoculated on agar medium in a petriplate at 4 equidistant points from the centre, where 7mm fungal agar disc was inoculated and then incubated in darkness at 27°C for 48-72 hrs. In parallel, PDA plate inoculated only with *R. solani* was kept as pure control. Percentage of growth inhibition (% GI) was calculated by comparing the radial distance of fungal growth towards each spot inoculation with ethanol control. IC_50_ was calculated by regression equation analysis.

### Phytotoxicity Assay

Phytotoxicity assay was conducted to ascertain the impact of the isolated bacterial endophyte on tomato seedlings’ health. Surface sterilized seeds were bio-primed with the pure culture of *B. siamensis*, with the microbial load adjusted to ≥10^8^ cfu/ml by diluting with sterile saline water. In contrast, uncoated surface-sterilized seeds were kept as control. Seeds were then kept for incubation with continuous agitation (150-200 rpm) at 27°C for 24 hrs. (Xia et al., 2015) After air drying, seeds were allowed to germinate on sterile filter paper immersed with autoclaved distilled water. After 9 days of incubation at 27°C, the seedlings were observed for the basic growth parameters such as germination percentage, hypocotyl length, root length, and seedlings’ wet weight.

### Data Analysis

All the experiments were conducted with 3 sets of replication. For germination assay, 20 seeds were used in each replication of 3 in square Petri plates (100mm diameter). For the alignment of the sequences, software Clustal W was used. The evolutionary history is inferred using the Neighbor-Joining method. A tree with 1000 bootstrap replicates was constructed using MEGA-X. Heatmap was produced through online software Heatmapper (www.heatmapper.ca).

## RESULTS

### Isolation, Identification, and Phylogenetic Analysis

Seventy-four bacterial endophytes were isolated from the various tissues of root, hypocotyl, and cotyledon of tomato plants of both the organic varieties (V1 and V2) using the culture-dependent technique. The majority of the isolates (59.4%) were obtained from the V1 variety. All the 74 isolates were grouped into 13 species using 16S rRNA based molecular identification. Comparing the two varieties, Pusa ruby (V1) harbored all the 13 species identified while the local variety (V2) possessed less diverse endophytic populations as only four species inhabited in it. All the bacterial isolates belonged to the phylum Firmicutes. The details of isolates concerning identification, accession number, similarity percentage, and source are summarized in **Table 1**. In the V1 variety, *Bacillus safensis* FO-36b (T) and *Bacillus siamensis* KCTC 13613(T) were the dominant species with relative abundance (RA) of 52.3 and 18.2%, respectively. In V2, *Bacillus australimaris* strain MCCC 1A05787 and *Bacillus safensis* strain NBRC 100820 were the dominant species with RA of 40 and 36.6%, respectively (**Figure 1**).

**TABLE 1.** Isolated endophytic *Bacillus* species from tomato seeds with their accession numbers.

| S.No. | Bacterial endophytic strains | Total No. of isolates | Source | % similarity of the sequence | Accession number |
| --- | --- | --- | --- | --- | --- |
| 1 | *Bacillus safensis* FO-36b | 2 | V1 | 99 | CP010405.1 |
| 2 | *Bacillus safensis* FO-36b(T) | 23 | V1 | 100 | ASJD01000027 |
| 3 | *Bacillus safensis* strain NBRC 100820 | 12 | V1,V2 | 99 | NR_113945.1 |
| 4 | *Bacillus australimaris* strain MCCC 1A05787 | 14 | V1,V2 | 99 | NR_148787.1 |
| 5 | *Bacillus australimaris* NH7I_1(T) | 1 | V1 | 100 | JX680098 |
| 6 | *Bacillus amyloliquefaciens* DSM7 | 1 | V1 | 99 | FN597644.1 |
| 7 | *Bacillus amyloliquefaciens* strain MPA 1034 | 2 | V1,V2 | 99 | NR_117946.1 |
| 8 | *Bacillus nakamurai* strain NRRL B-41091 | 1 | V1 | 99 | NR_151897.1 |
| 9 | *Bacillus siamensis* KCTC 13613(T) | 8 | V1 | 100 | AJVF01000043 |
| 10 | *Bacillus zhangzhouensis* strain MCCC 1A08372 | 7 | V1,V2 | 99 | NR_148786.1 |
| 11 | *Bacillus zhangzhouensis* DW5-4(T) | 1 | V1 | 99.91 | JOTP01000061 |
| 12 | *Bacillus subtilis* subsp. *inaquosorum* strain KCTC 13429 | 1 | V1 | 96 | CP029465.1 |
| 13 | *Bacillus subtilis* subsp. *subtilis* strain 168 | 1 | V1 | 99 | NR_102783.2 |

**FIGURE 1.**
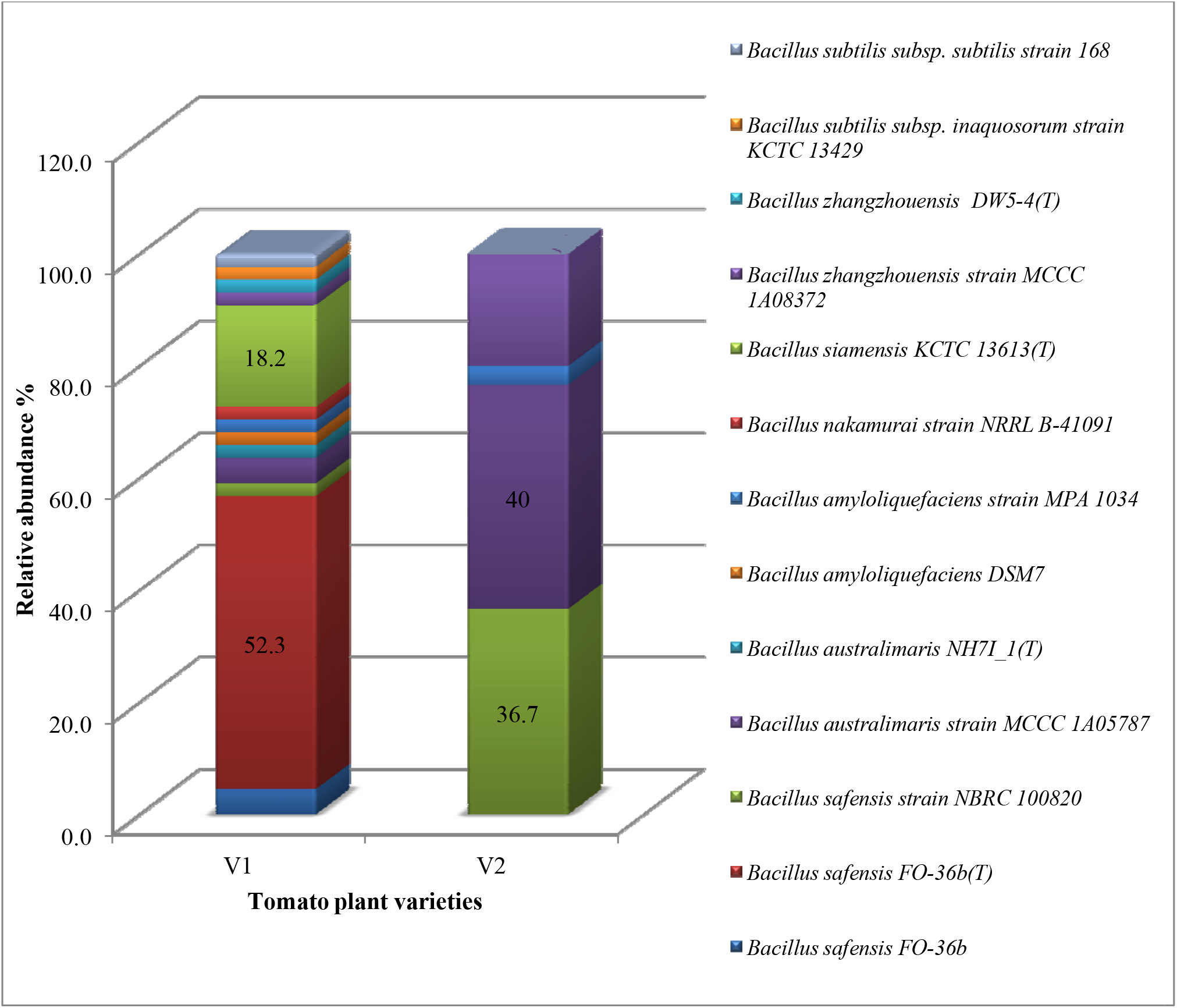
Taxonomic profiles of the bacterial community in each system at the species level with the relative abundance

In V1 isolates, only two endophytic bacterial strains, namely *Bacillus safensis* FO-36b(T) and *Bacillus siamensis* KCTC 13613(T), were isolated from all the three parts of tomato seedling (root, hypocotyl, and cotyledon), and other bacterial endophytic species were only exclusive to one or two tissues. However, three out of four species isolated from the V2 variety, namely, *Bacillus safensis* strain NBRC 100820, *Bacillus australimaris* strain MCCC 1A05787, *Bacillus zhangzhouensis* strain MCCC 1A08372, were found to inhabit the three parts of tomato seedling (root, hypocotyl, and cotyledon), whereas, *Bacillus amyloliquefaciens* strain MPA 1034 was found only in root region. The evolutionary history was inferred using the Neighbor-Joining method (**Figure 2**). The Phylogenetic analysis showed an evolutionary relationship between the isolated strains and all the species grouped into two broad categories. *B. siamensis* did not group with any other species identified by us.

**FIGURE 2.**
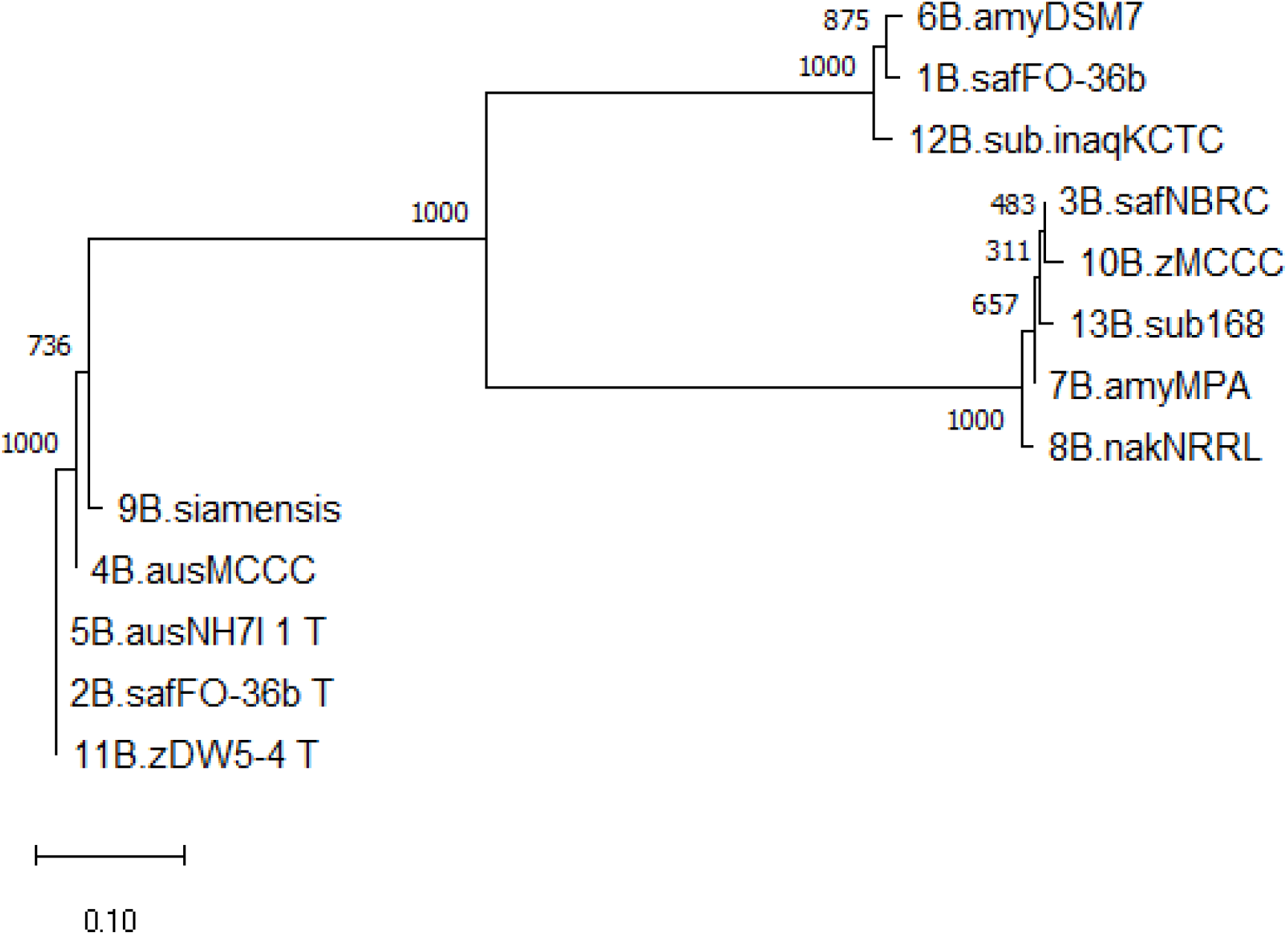
Phylogenetic tree constructed using 16S rRNA gene sequences of 13 different strains of *Bacillus* species and bootstrap values are indicated at the nodes

### Distribution, Diversity, and Richness of Endophytic Bacterial Isolates

Diversity indices were calculated between the bacterial endophytes isolated from each tissue of the two varieties of tomato plants used in the study (**Table 2**). Shannon diversity (H’) was maximum in the hypocotyl (1.45) and cotyledon (1.33) of V1 variety, followed by the root (1.30). Least diversity was reported in the cotyledon region of V2 variety (0.92). Simpson’s index of diversity was maximum in the root (1) of V2, followed by cotyledon (0.9) of V1. Species richness was determined by counting the number of species in each group and was found maximum in hypocotyl (n = 8) of V1 variety followed by the root (n = 5) and cotyledon (n = 4) of V1. Magalef’ index, calculated to estimate the evenness between the species of both the types, was found to be highest in hypocotyl (2.1) of V1variety. Species shared between V1 and V2 were highest in hypocotyl, resulting in a high value of Sorenson’s similarity index (0.266) (**Table 3**). A Venn diagram illustrated the species’ number and the relationship between the isolated species within the same variety (**Figure 3**). Interestingly, the V1 variety of tomato (Pusa Ruby) contains a more diverse population of endophytic bacteria as compared to V2 (**Table 2**).

**TABLE 2.** Diversity indices of bacterial endophytes isolated from V1 and V2 variety of tomato.

|  | V1 |  |  | V2 |  |  |
| --- | --- | --- | --- | --- | --- | --- |
|  | Hypocotyl | Root | Cotyledon | Hypocotyl | Root | Cotyledon |
| **Shannon diversity** | 1.45 | 1.3 | 1.33 | 0.94 | 1.28 | 0.92 |
| **Simpson's Index** | 0.32 | 0.29 | 0.1 | 0.36 | 0 | 0.38 |
| **Simpson's index of diversity** | 0.67 | 0.7 | 0.9 | 0.63 | 1 | 0.61 |
| **Magalef' index** | 2.1 | 1.67 | 1.87 | 0.91 | 1.3 | 0.83 |
| **Evenness** | 0.69 | 0.8 | 0.96 | 0.85 | 0.92 | 0.84 |
| **Species Richness** | 8 | 5 | 4 | 3 | 4 | 3 |

**TABLE 3.** Comparison of different similarity indices among different regions of two organic varieties of tomato.

| **V1 vs. V2** | **Species shared** | **Jaccard's SI** | **Sorensen's SI** |
| --- | --- | --- | --- |
| **Root** | 1 | 0.1 | 0.182 |
| **Hypocotyl** | 2 | 0.153 | 0.266 |
| **Cotyledon** | 0 | 0 | 0 |

**FIGURE 3.**
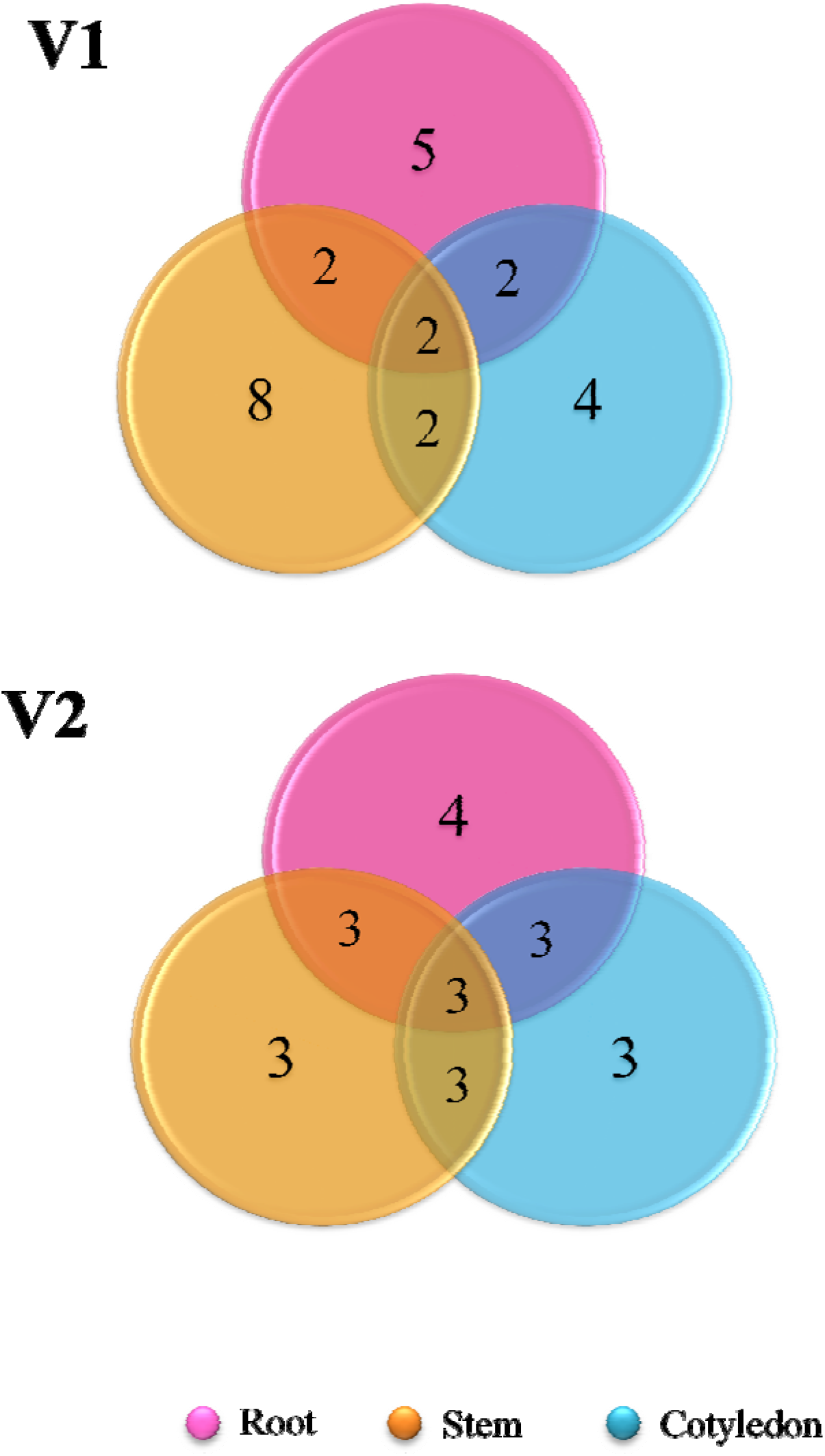
Venn diagram representing the shared species of isolated bacterial endophytes within the variety

### Antifungal Activity of the Isolated Endophytic Bacteria

All the bacterial endophytes isolated from the two organic tomato varieties were screened for their antifungal activity against five economically important fungal pathogens of tomato crop viz. *R. solani, V. lateritium, B. cinerea, A. solani*, and *F. solani* through dual culture assay (**Figure 4**). The dual culture bioassay’s key purpose was based on a bio-prospecting strategy to select potential endophytes with having antifungal activity. Among all the isolates, *B. siamensis* KCTC 13613(T) exhibited the highest antifungal activity having percentage growth inhibition values ranging from 75 - 90%, against all the five major pathogens of the tomato crop (**Figure 5; Supplementary Table S1**). To the best of our knowledge, this is the first report on the antifungal activity of endophytic bacteria isolated from the organic varieties of tomato.

**FIGURE 4.**
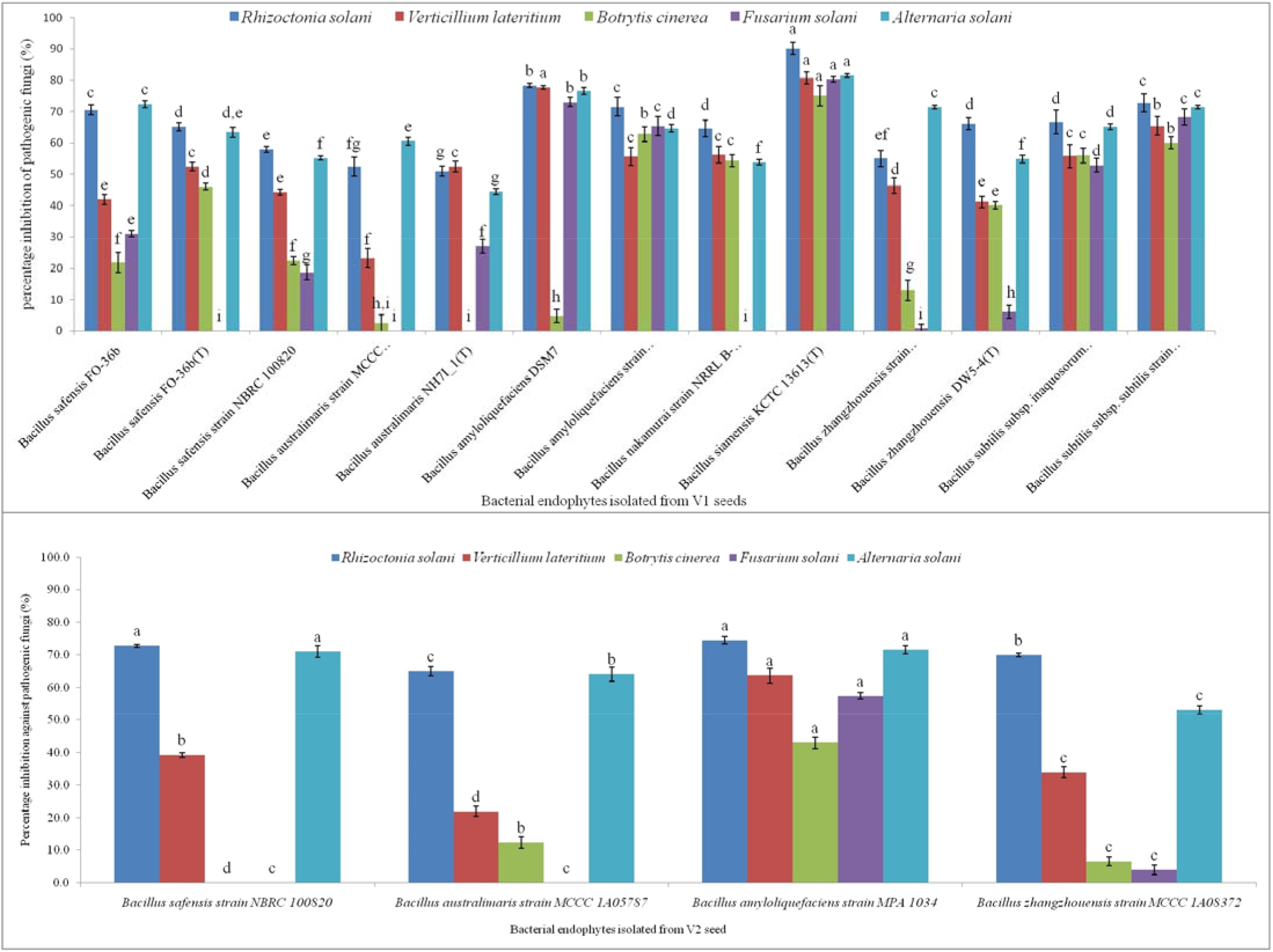
Antagonistic effect against five pathogenic test fungi by (A) *Bacillus* strains isolated from V1; (B) *Bacillus* strains isolated from V2

**FIGURE 5.**
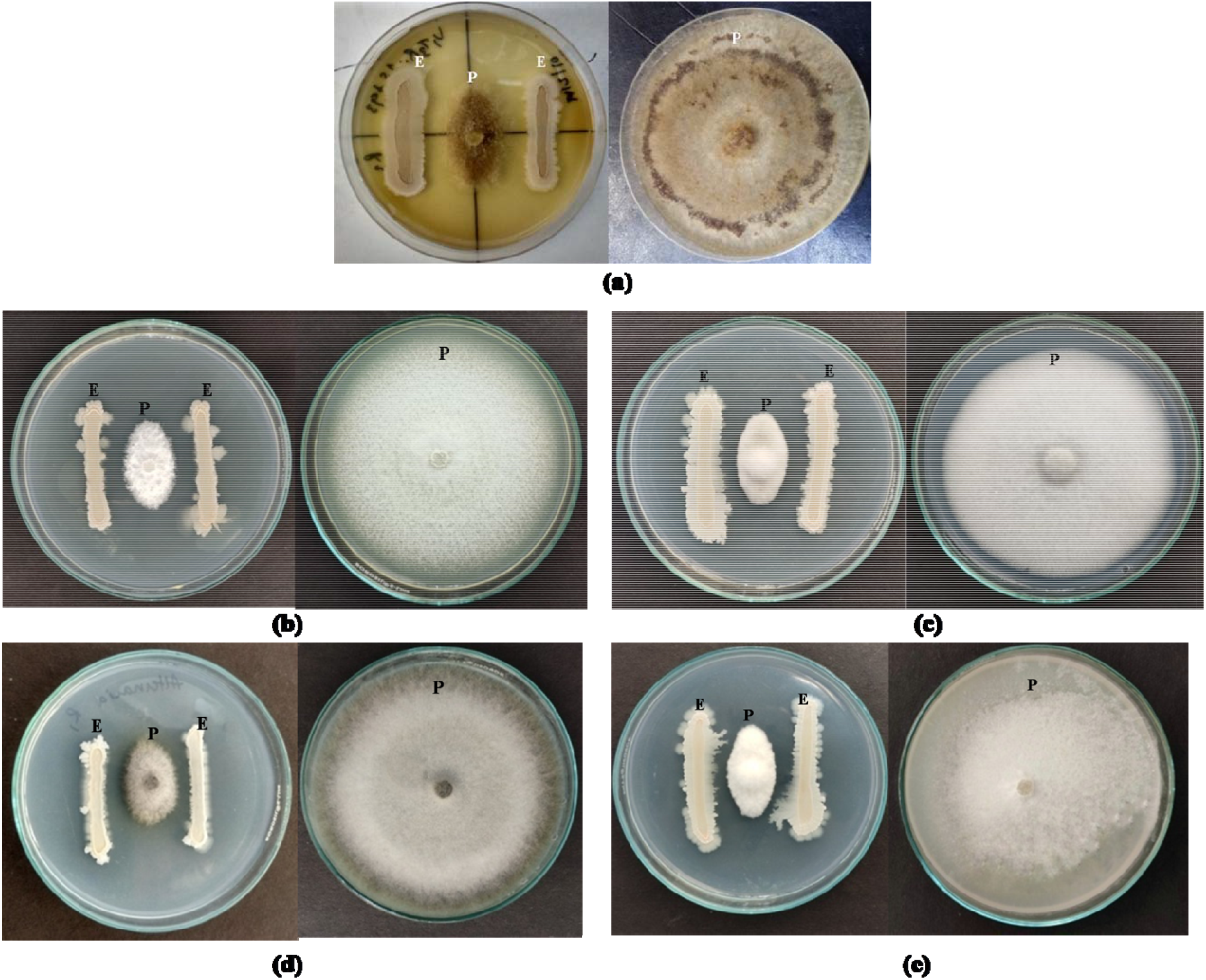
Antagonizing effect of *Bacillus siamensis* KCTC 13613(T) against (a) *Rhizoctonia solani*, (b) *Verticillium lateritium* (c) *Botrytis cinerea*, (d) *Alternaria solani*, and (e) *Fusarium solani* after six days of inoculation (‘E’ represents endophytic bacterial strain whereas ‘P’ represents Pathogenic fungi)

*B. amyloliquefaciens* was found to be the next best species. The activity pattern of *B. safensis* varied from strain to strain. The most active strain of *B. safensis viz. Bs safensis* strain NBRC 100820 isolated from variety V2 recorded >70% growth inhibition activity against *R. solani* and *A. solani. Simultaneously, B. australimaris, B. nakamurai*, and *B. zhangzhouensis* showed very low to nil activity against the selected pathogens (**Supplementary Table S2**). Heatmap dendrogram revealed that the antifungal activity of the tested strains against *R. solani* positively correlated with *A. solani* while activity against *B. cinerea* correlated with activity against *F. solani* (**Figure 6**).

**FIGURE 6.**
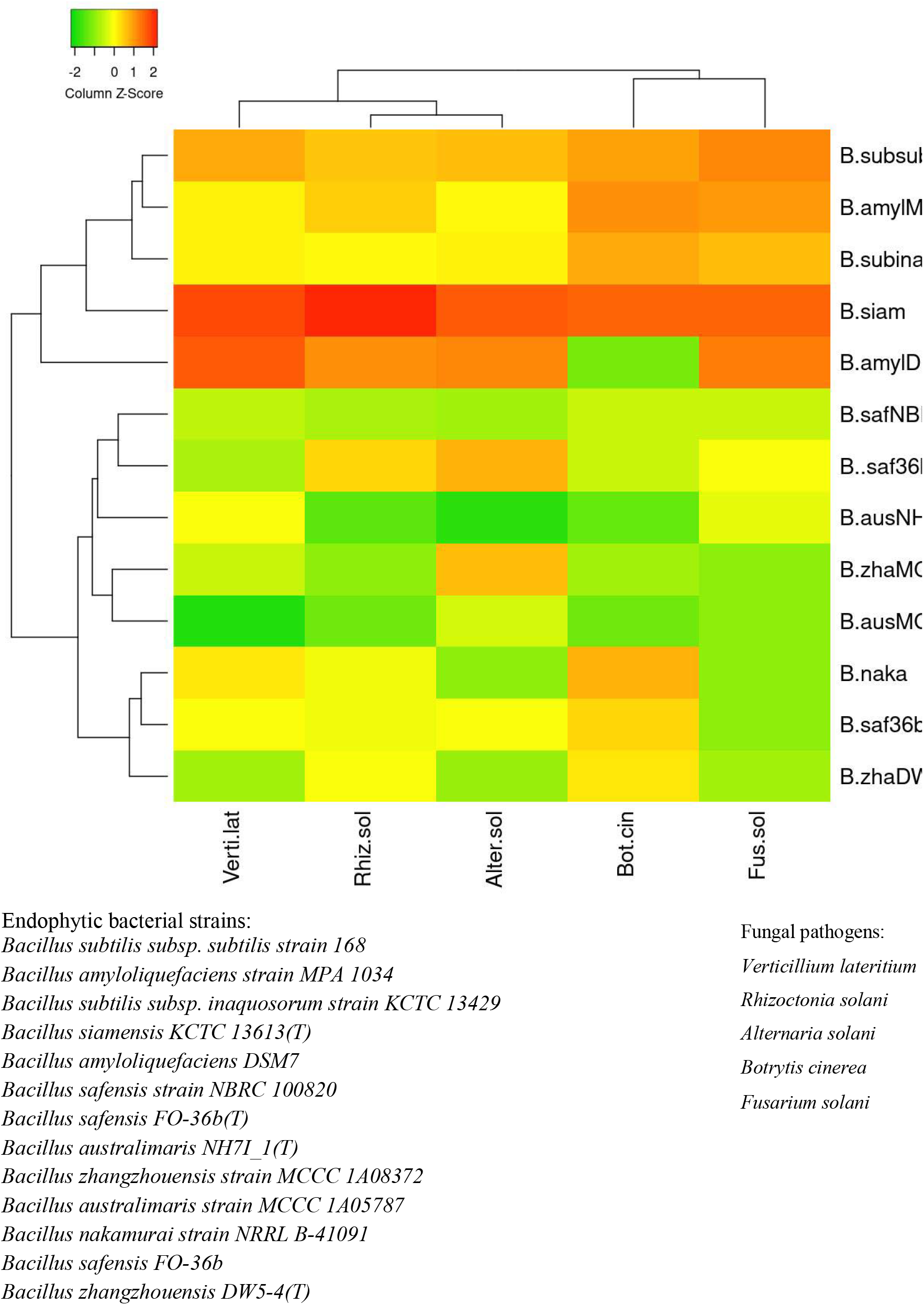
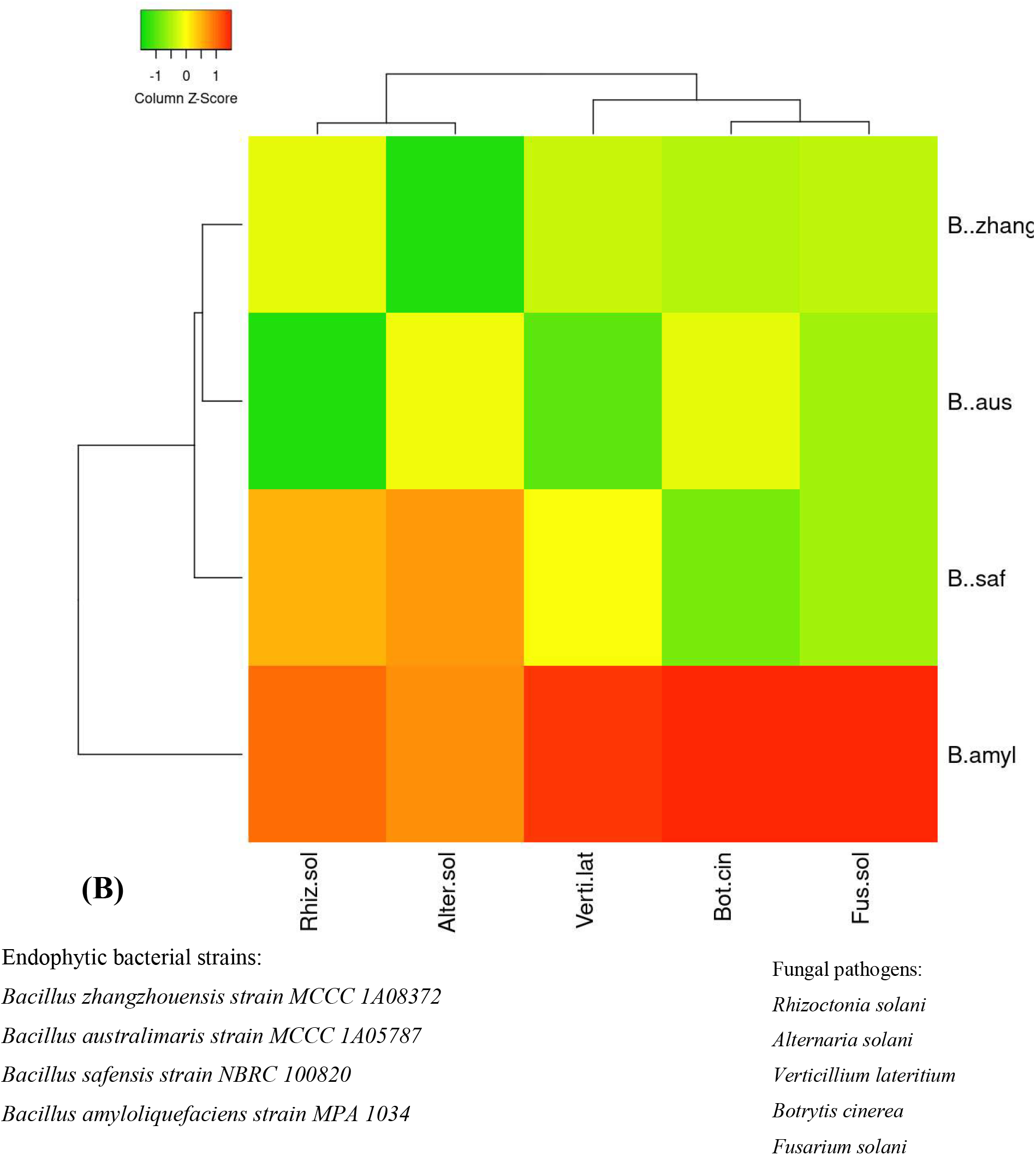
Heat map illustrating the strength of antifungal activity of different *Bacillus* strains of (A) V1 and (B) V2 against all five pathogenic fungi with respect to other strains

The endophytic population from variety V1 has been observed more antagonistic against all the five pathogenic fungi than the V2. None of the endophytes found active against all the five test pathogens. More than 95% of endophytic bacteria of V1 suppressed the growth of *R. solani* in dual culture assay with antagonistic activity up to 90%. Meanwhile, 17% of its population showed the antagonistic effect against all the test pathogens with over 70% inhibition.

### Antifungal Activity of Lipopeptide

Ethanol extract of lipopeptide obtained from the culture of *B. siamensis* was subjected to bioassay to examine its antifungal activity against *R. solani*. Dose response was observed with R^2^ value of 0.99 88.8 % growth inhibition of *R. solani* was observed at 300 ppm extract of *B. siamensis* (**Figure 9**). The IC_50_ value of 72 ppm was obtained using regression equation (**Supplementary Figure S1).**

### Lipopeptide Profiling by UPLC-HDMS

To identify the compound responsible for the antifungal activity in *B. siamensis*, lipopeptide extraction was done from *B. siamensis* culture. Chromatographic separation and Mass Spectrometry of ethanol extract of the lipopeptide was performed on UPLC-H class with Synapt G2-Si-High Definition Mass Spectrometry (HDMS) system equipped with an auto sampler. **Figure 7** reveals the mass spectrum of the analyte showing the presence of the molecular peaks at m/z 994.8, 1008.77, 1022.72, 1036.74, 1050.75, 1064.77, 1096.86, 1045.77, 1059.79, and 1079.81. These masses were assigned to Surfactin and Bacillomycin D lipopeptides (**Table 4**). The general molecular structures of the isolated antifungal lipopeptides are presented in **Figure 8**.

**TABLE 4.** Main mass peaks of the lipopeptides produced by *Bacillus siamensis* mass spectrometry.

| Mass peaks (m/z) | Assignment |
| --- | --- |
| 994.8 | C12 Surfactin[M + H<sup>+</sup>]<sup>+</sup> |
| 1008.77 | C13 Surfactin[M + H<sup>+</sup>]<sup>+</sup> |
| 1022.72 | C14 Surfactin [M + H<sup>+</sup>]<sup>+</sup> |
| 1036.74 | C15 Surfactin [M + H<sup>+</sup>]<sup>+</sup> |
| 1050.75 | C16 Surfactin [M + H<sup>+</sup>]<sup>+</sup> |
| 1064.77 | C17 Surfactin [M + H<sup>+</sup>]<sup>+</sup> |
| 1096.86 | Linear C18 Surfactin |
| 1045.77 | C15 Bacillomycin D [M + H<sup>+</sup>]<sup>+</sup> |

**FIGURE 7.**
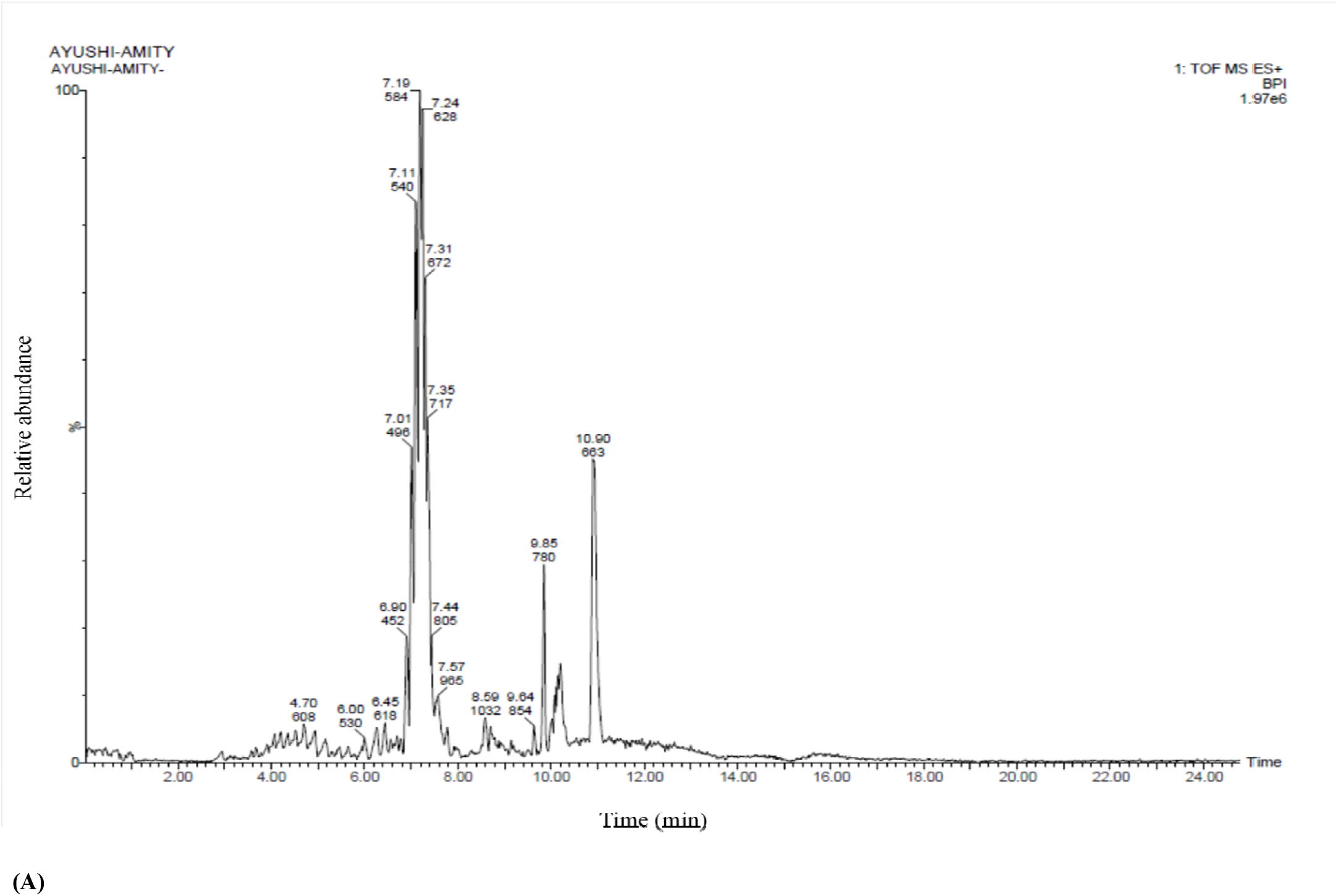
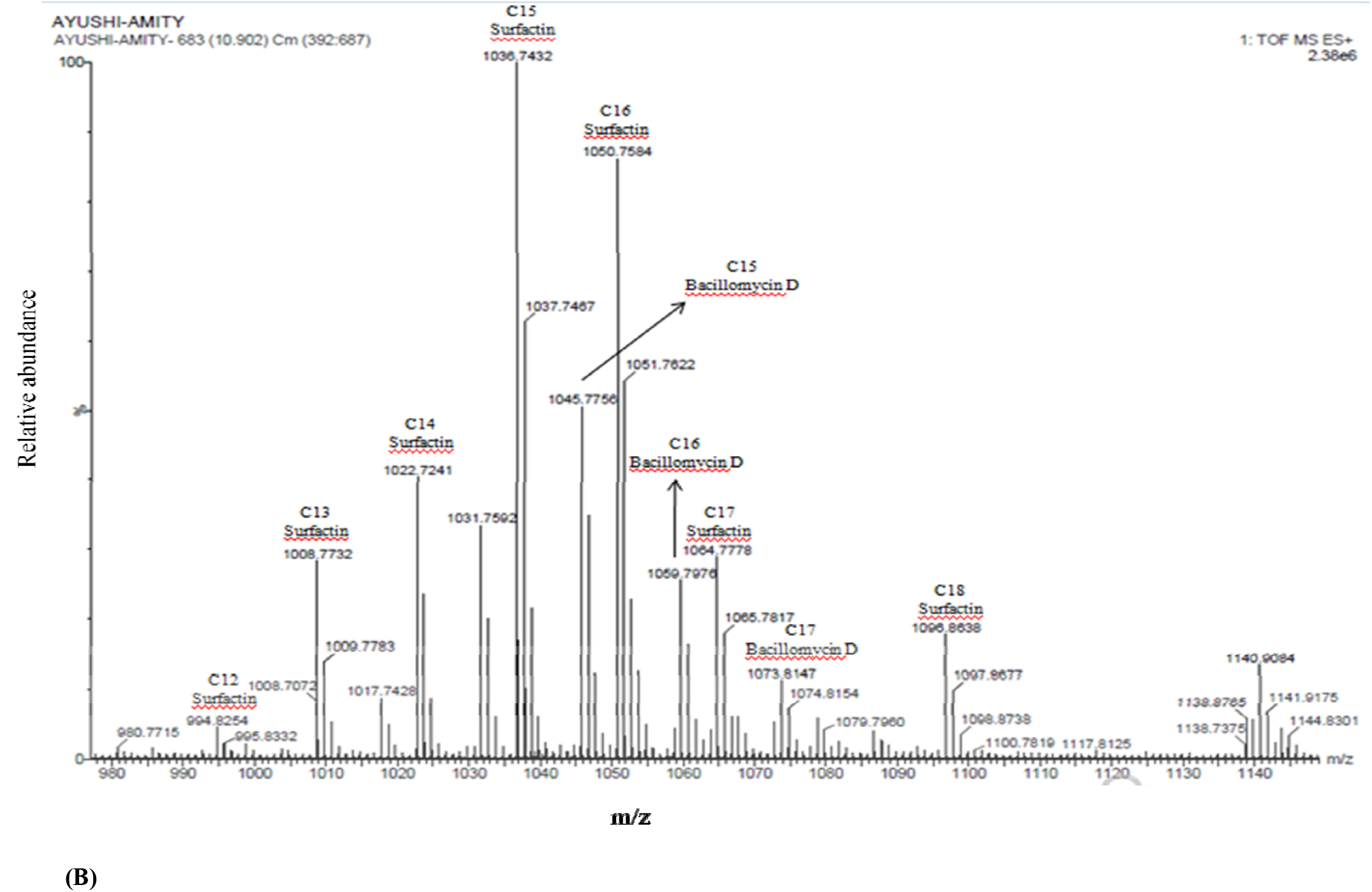
A) UPLC chromatogram of lipopeptides extracted from *Bacillus siamensis* strain; B) HDMS accurate mass revealed the production of Surfactin and Bacillomycin D analogues

**FIGURE 8.**
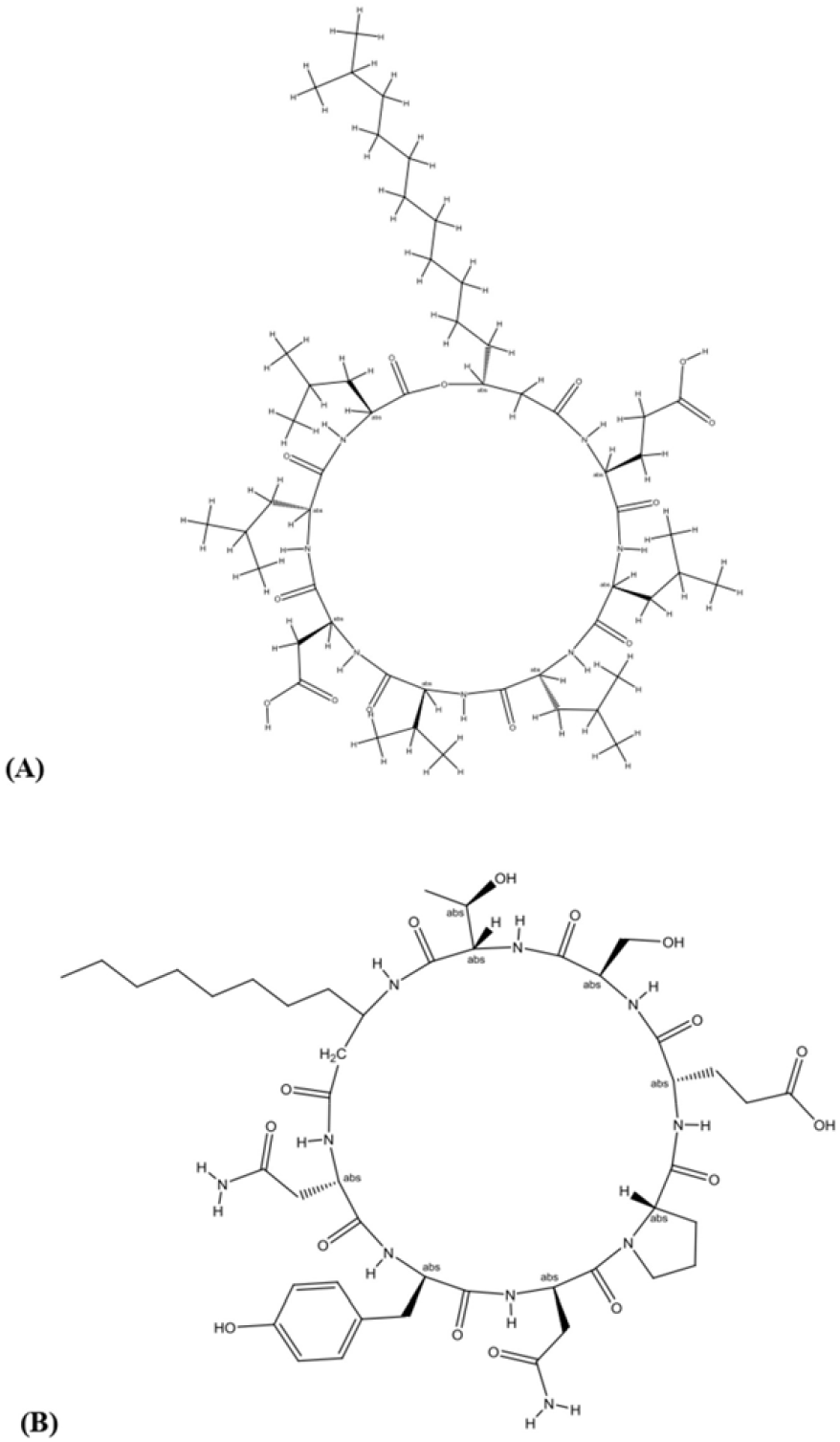
General molecular structure of lipopeptides (A) Surfactin and (B) Bacillomycin isolated from *B. siamensis*

**FIGURE 9.**
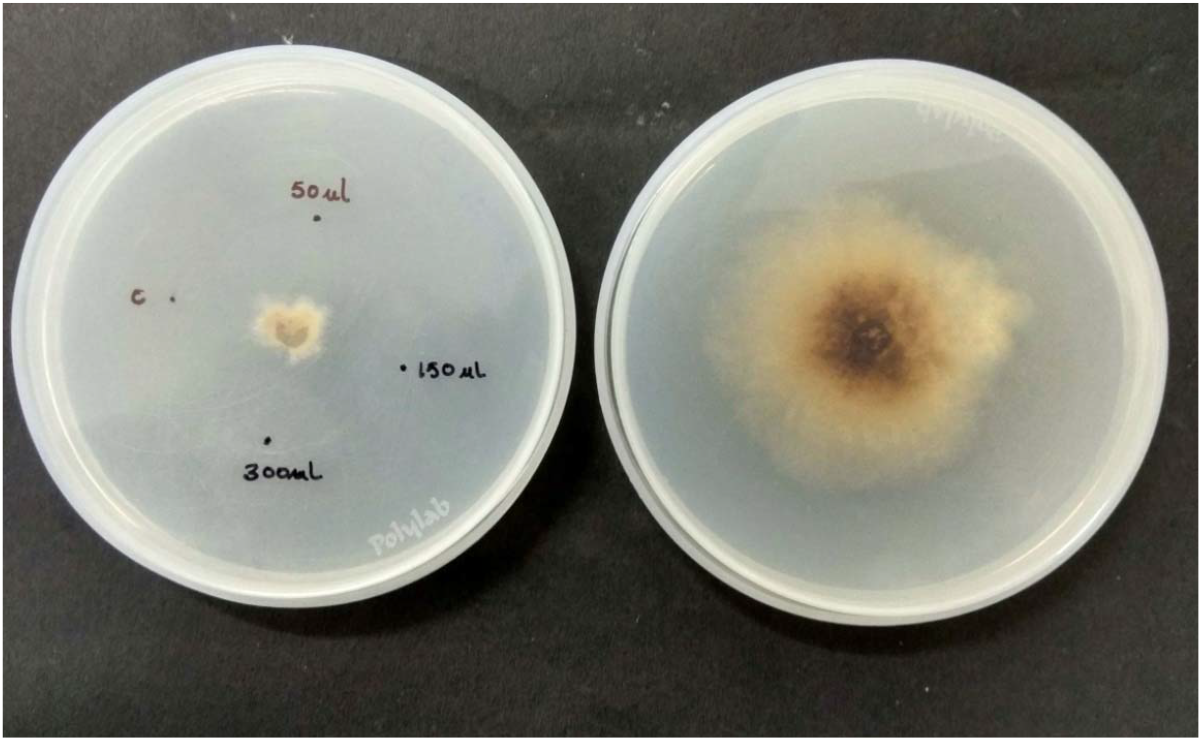
Antifungal bioassay of lipopeptide extracted from *Bacillus siamensis* strain at three different concentrations (A) 50 ppm, 150 ppm and 300 ppm; (B) Pure control

### Phytotoxicity Assay

To assess whether *B. siamensis* has any detrimental effect on plant growth, a phytotoxic assay was performed by seed bacterization of tomato seeds (**Figure 10**). It was observed that the treatment with the pure culture of this strain did not hamper the germination and seedling growth (**Table 5**). Instead, it increased the fresh biomass of tomato seeding by 41.6%, hypocotyl length by 32.9%, root length by 49.1%, besides a 6.7% increase in germination.

**TABLE 5.**
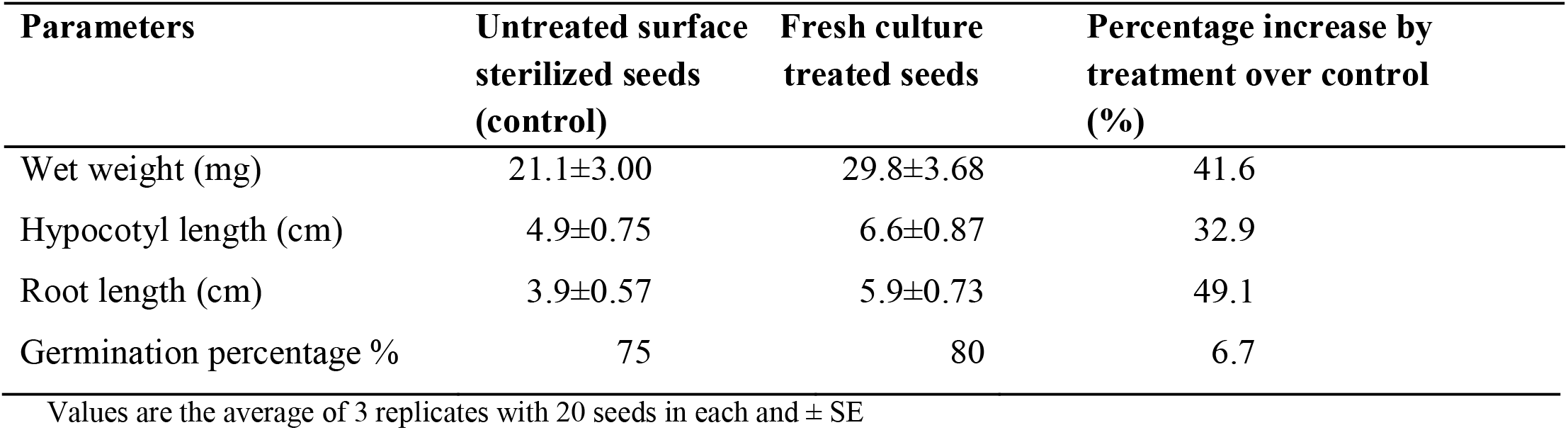
Growth study of seeds primed with the pure culture of *B. siamensis* with respect to the control.

**FIGURE 10.**
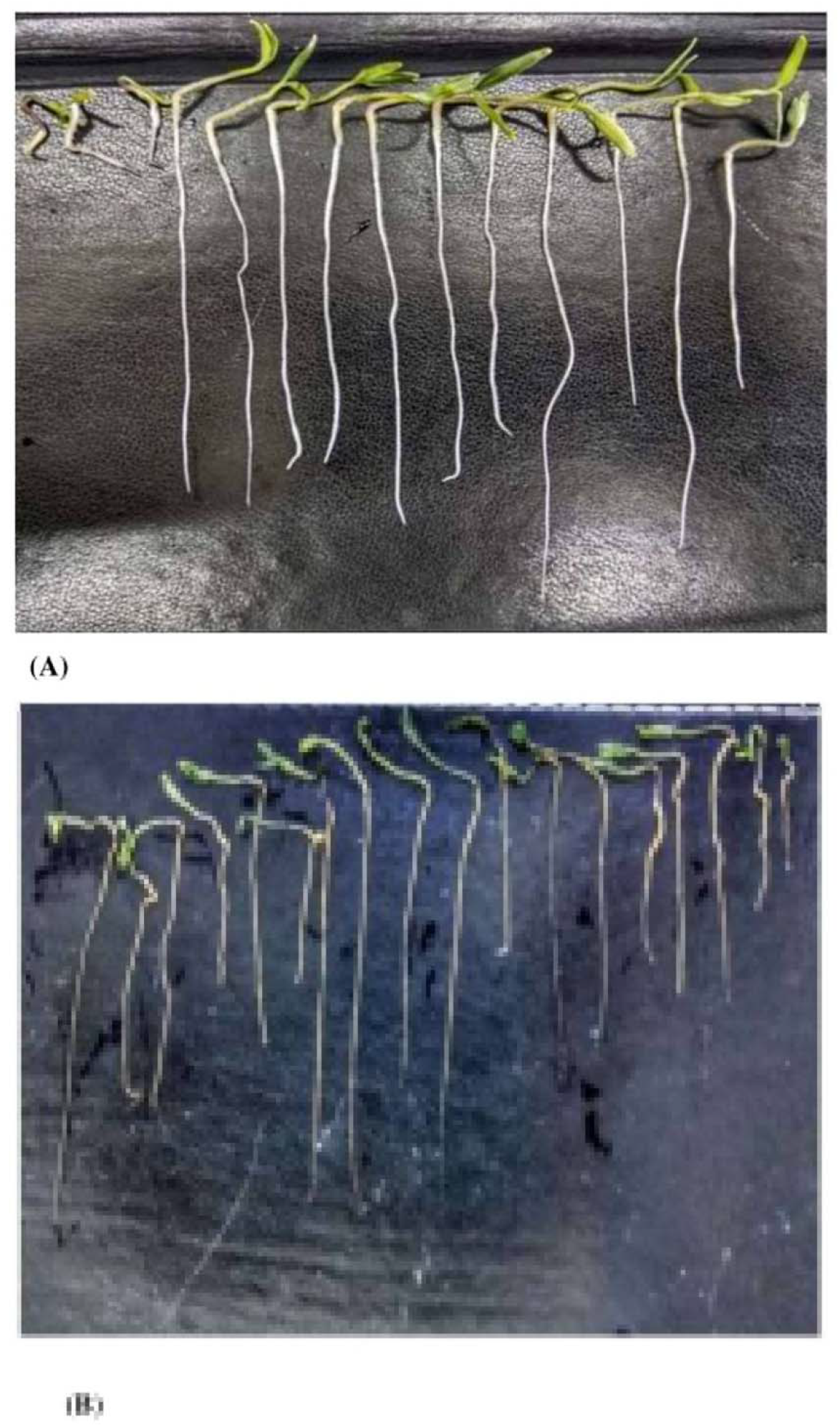
Germinated seedlings from bio-primed tomato seeds after 9 days (A) and control (B)

## DISCUSSION

The present research covers two organic tomato varieties for greater cultivable diversity of endophytic bacteria and their antifungal ability against selected fungal pathogens. The seedlings of the V1 tomato plant variety were found to be rich in species abundances and the diversity of bacterial endophytes. Ours is the first report on the diversity study of endophytic bacteria from organic tomato plants. The plausible reason for the disparity in endophyte diversity between the two tomato plant varieties could be the variations in the rhizospheric microbiome that probably contribute to differential bacterial colonization in the plant endosphere (Liu et al., 2017; Compant et al., 2010). Species richness was found maximum in the hypocotyl of the seedling (n=8) of V1. These findings indicate that endophytic bacteria can exhibit a tissue-specific distribution, which has also been reported from other systems (Reinhold-Hurek and Hurek, 2011; Thomas and Reddy, 2013; Xia et al., 2015). Previous studies have shown the species specificity of endophytes. The difference in endophytic assemblies in different tissue types can be due to the difference in their potential to use the substrate (Huang et al., 2008; Chowdhary and Kaushik, 2015). Yang et al. (2011) reported 72 bacterial endophytes, including 45 from the stem and 27 from the healthy tomato plant leaves, and found *Brevibacillus brevis* W4, an endophyte antagonistic to *B. cinerea*. We believe that different agro-climatic locations (V1 from Maharashtra and V2 from Andhra Pradesh) resulted in endophytic population variations in the current study. The cultivable bacteria obtained from the tomato varieties’ seedlings were similar to the phyla found inside the seedlings. This suggests that tomato seeds may contain a specific subset of bacteria that are likely to reach seed during the reproductive phase. These bacteria are most likely to play different roles in seed health seedling growth (Lopez et al., 2018). The host genotype is reported to play an essential role in managing the associated plant microorganisms, particularly the endophytes (Lundberg et al., 2012; Podolich et al., 2015; Upreti and Thomas, 2015). Also, there are indications of endophytic bacterial transmission via seeds, which might clarify their possible integral interaction with a specific host varietal (Truyens et al., 2014).

Despite being identical in the presence of species, our findings show that under-regulated conditions, not all bacteria inhibit mycelial growth; however, they vary in their ability to synthesize other inhibitory molecules. In comparison to the endophytes in variety V2, the V1 endophytic population is increasingly antagonistic to all five test fungi. The most potent antagonistic endophyte was identified through 16S rRNA sequencing as *B. siamensis* KCTC 13613(T). There was no physical contact between the isolates and the pathogen in the inhibition zone, indicating that the isolated active *Bacillus* species may generate definite antifungal substances that impede the mycelial growth (Lee et al., 2008). *B. siamensis* KCTC 13613(T) exhibits more antagonistic activity than other species against all the selected fungal pathogens. The z-score clustering facilitates the bacterial species relationship between the isolates in relation to the fungal pathogens. A higher z value suggests that genotypes will be better clustered by function, suggesting a clustering result, which is more biologically important (Bhattacharya et al., 2012). In variety, V1 *B. cinerea and F. solani* are less susceptible to antifungal behavior of some endophytic species or have similar responses to most of the bacterial species. Likewise, *R. solani* and *A. solani* linked similar responses with those of *V*. *lateritium*. This clustering is not by chance but because of a computer program that aims to close similar things together. However, in variety V2), *B. amyloliquefaciens strain* MPA 1034 is notable for maximum antifungal activity against all pathogen fungi.

Many endophytic and non-endophytic *Bacillus* spp. including *B. siamensis*, have been reported to produce a wide variety of structurally different antagonistic substances through secondary metabolism (Fira et al., 2018). Interestingly, the strains producing non-ribosomally synthesized lipopeptides and peptides have shown enhanced fungicidal activities (Dimkic et al., 2013; Etchegaray et al., 2008). The LC-MS/MS-based analysis of the extract further confirmed the product of surfactin derivatives, iturin, and fengycin by *Bacillus* sp. (Jasim et al., 2016). This is the first experimental evidence of the presence of these antifungal lipopeptides in *B. siamensis*. The IC_50_ value of 72 ppm showed the high potency of the crude extract obtained from the pure culture of *B. siamensis* to inhibit the growth of *R. solani*, thus further confirming that the antifungal activity of the *B. siamensis* is due to lipopeptides. Earlier, it was predicted through genome sequencing that *B. siamensis* contains Sufactin and Bacillomycin D genes (Pan et al., 2019). However, it is for the first time that it has been extracted and confirmed in the culture broth. It is believed that bioactive compounds producing bacterial endophytes can be an effective biological agent and a powerful tool for the development of a formulation against fungal pathogens in crop protection and for promoting plant growth. The mechanism of action of lipopeptides might depend on the structural and functional properties of lipopeptides (Zhang et al., 2013). Bacillomycin L antifungal activity against *R. solani Kühn*, which includes a specific association with intact fungal hyphae, has been extensively investigated using different fluorescent methods, gel retardation experiments, and electron microscopy (Zhang et al., 2013).

The majority of *B. siamensis* strain isolation has been reported from rhizosphere or other sources other than endophytic (Yoo et al., 2020; Hussain and Khan. 2020; Islam et al., 2019; Pastor-Bueis et al., 2017). Antifungal activity of filtrate obtained from the culture of *B. siamensis* has been previously reported, such as in a study by Putri et al., 2020, ethyl acetate extract of fermentation filtrate of *B. siamensis* showed antifungal activity against *Aspergillus niger*. Various *Bacillus* strains, including *B. siamensis*, has been identified to produce biosurfactants as surfactin variants based on analytical methods and surfactin gene phylogenetic analysis (Mehetre et al., 2019). A similar study conducted by Pan et al. (2019) reported *B. siamensis* to produce sets of bacillibactins, fengycins, bacillomycins, and surfactins through the mining of genome and metabolic profiling. The PCR study demonstrated the existence of genes (i.e., surfactin synthetase D and bacillomycin synthetase D) involved in cyclic lipopeptide biosynthesis against multidrug-resistant aquatic bacterial pathogens (Xu et al., 2018). This concludes that so far, no endophytic strain of *B. siamensis* with antifungal potential has been reported to produce surfactin and Bacillomycin D. Complete isolation and identification of these lipopeptides from *B. siamensis* KCTC 13613(T) isolated from the varieties of tomato plants is first to be reported. This indicates that the use of beneficial bacteria native to their host plants may increase the success rate in screening bio-control experiments because these microbes are likely to be better adapted to their host and its associated environmental conditions than are strains retrieved from culture collections (Karimi et al., 2016; Köbrel et al., 2013).

A study by Karthik et al. (2017), compared to uninoculated control, bacterial inoculation treatment with endophytic strains on tomato seeds, significantly improved seed germination, seedling growth, vigor index, and biomass production. *Rhizobium taibaishanense* (RBEB2), *Pseudomonas psychrotolerance* (REB4) and *Microbacterium testaceum* (RBEB1) had significant positive effects on the germination of tomato seeds and vigor index. *Bacillus subtilis* (RBEB6) enhanced the biomass as well as root and shoot length of tomato seedling.

The data presented here collectively support the notion that soil properties and rhizospheric microflora can affect the endophytic microflora. To the best of our knowledge, this is the first report of the isolation and diversification of bacterial endophytes from organic tomato seeds; however, we only found the presence of *Bacillus* species. Comparatively, Pusa Ruby has a more diverse and biologically active endophytic population of bacteria, and lipopeptide producing *B. siamensis* is a promising antifungal bio-control agent.

## AUTHOR CONTRIBUTIONS

NK and ND conceived the idea, secured funding, planned the work and guided first author. AS (first author) performed the experimental work and wrote the manuscript. AS (third author) helped in data analysis. NK, AS (third author), and ND read and reviewed the manuscript. AB, MS and YS conducted the molecular identification of all the bacterial isolates. TM supported in lipopeptide extraction process.

## ACKNOWLEDGMENT

The authors are thankful to the Amity University Uttar Pradesh, Noida (India) for providing the infrastructure for the smooth conduction of experiments.

