## Supplementary Table S1 for "Screening of tomato seed bacterial endophytes for antifungal activity reveals lipopeptide producing *Bacillus siamensis* as a potential bio-control agent": Supplementary_Table_S1.docx

**Supplementary Table S1**: Inhibition percentage of isolated bacterial endophytes from V1 variety against plant pathogenic fungi

| **Bacterial endophytes** | **Percentage inhibition of fungal pathogens (%)** | | | | |
| --- | --- | --- | --- | --- | --- |
|  | ***Rhizoctonia solani*** | ***Verticillium lateritium*** | ***Botrytis cinerea*** | ***Fusarium solani*** | ***Alternaria solani*** |
| *Bacillus safensis FO-36b* | 70.5^c^ | 41.9^e^ | 21.8^f^ | 31.0^e^ | 72.4^c^ |
| *Bacillus safensis FO-36b(T)* | 65.2^d^ | 52.5^c^ | 46.0^d^ | 0.0^i^ | 63.4^d,e^ |
| *Bacillus safensis strain NBRC 100820* | 57.9^e^ | 44.3^e^ | 22.4^f^ | 18.7^g^ | 55.2^f^ |
| *Bacillus australimaris strain MCCC 1A05787* | 52.5^f,g^ | 23.2^f^ | 2.5^h,i^ | 0.0^i^ | 60.6^e^ |
| *Bacillus australimaris NH7I_1(T)* | 51.0^g^ | 52.4^c^ | 0.0^i^ | 27.1^f^ | 44.5^g^ |
| *Bacillus amyloliquefaciens DSM7* | 78.3^b^ | 77.6^a^ | 4.8^h^ | 73.0^b^ | 76.6^b^ |
| *Bacillus amyloliquefaciens strain MPA 1034* | 71.5^c^ | 55.7^c^ | 62.8^b^ | 65.4^c^ | 64.6^d^ |
| *Bacillus nakamurai strain NRRL B-41091* | 64.6^d^ | 56.3^c^ | 54.3^c^ | 0.0^i^ | 53.8^f^ |
| *Bacillus siamensis KCTC 13613(T)* | 90.1^a^ | 80.7^a^ | 75.1^a^ | 80.3^a^ | 81.5^a^ |
| *Bacillus zhangzhouensis strain MCCC 1A08372* | 55.0^e,f^ | 46.3^d^ | 12.9^g^ | 0.8^i^ | 71.4^c^ |
| *Bacillus zhangzhouensis DW5-4(T)* | 66.1^d^ | 41.2^e^ | 40.1^e^ | 6.1^h^ | 54.8^f^ |
| *Bacillus subtilis subsp. inaquosorum strain KCTC 13429* | 66.6^d^ | 55.8^c^ | 56.0^c^ | 52.8^d^ | 65.2^d^ |
| *Bacillus subtilis subsp. subtilis strain 168* | 72.7^c^ | 65.4^b^ | 60.0^b^ | 68.3^c^ | 71.4^c^ |

Values are mean of 3 replications.

Means followed by a common letter are not significantly different at 5% level by DMRT.

‘a’ signifies highest value followed by ‘b’> ‘c’ > ‘d’.
