## Supplementary Table S2 for "Screening of tomato seed bacterial endophytes for antifungal activity reveals lipopeptide producing *Bacillus siamensis* as a potential bio-control agent": Supplementary_Table_S2.docx

**Supplementary Table S2:** Inhibition percentage of isolated bacterial endophytes from V2 variety against plant pathogenic fungi

| **Bacterial endophytic strains** | **Percentage inhibition of fungal pathogens (%)** | | | | |
| --- | --- | --- | --- | --- | --- |
|  | ***Rhizoctonia solani*** | ***Verticillium lateritium*** | ***Botrytis cinerea*** | ***Fusarium solani*** | ***Alternaria solani*** |
| *Bacillus safensis strain NBRC 100820* | 72.7^a^ | 39.2^b^ | 0.0^d^ | 0.0^c^ | 71.0^a^ |
| *Bacillus australimaris strain MCCC 1A05787* | 64.9^c^ | 21.9^d^ | 12.3^b^ | 0.0^c^ | 64.0^b^ |
| *Bacillus amyloliquefaciens strain MPA 1034* | 74.5^a^ | 63.6^a^ | 43.0^a^ | 57.4^a^ | 71.5^a^ |
| *Bacillus zhangzhouensis strain MCCC 1A08372* | 69.8^b^ | 33.9^c^ | 6.6^c^ | 3.9^c^ | 53.1^c^ |

Values are mean of 3 replications.

Means followed by a common letter are not significantly different at 5% level by DMRT.

‘a’ signifies highest value followed by ‘b’> ‘c’ > ‘d’.
