## Supplementary figures and images for "Screening of tomato seed bacterial endophytes for antifungal activity reveals lipopeptide producing *Bacillus siamensis* as a potential bio-control agent"

### Supplementary Figure S1.jpg

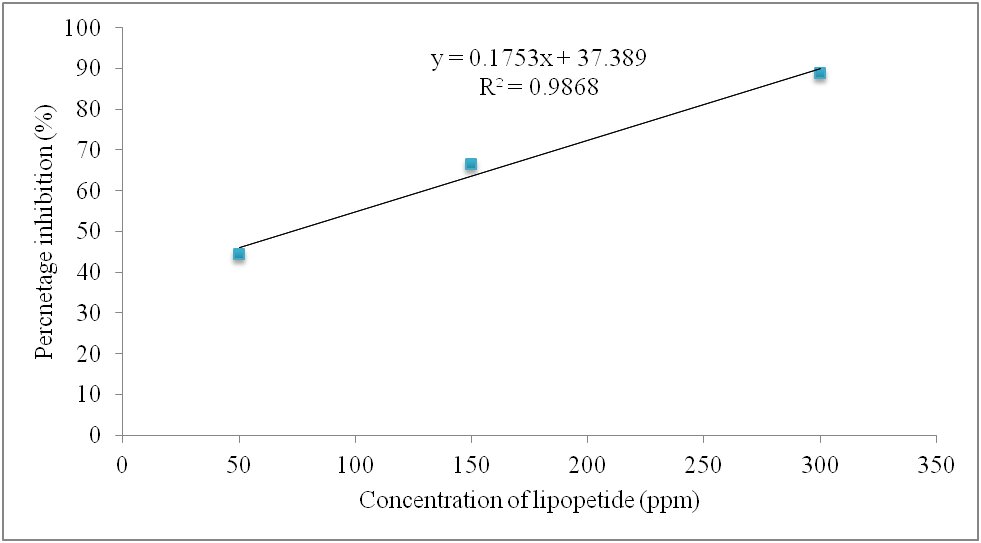
